# *Hylocereus polyrhizus* peel extract restricts DMBA-Croton oil induced skin carcinogenesis: An integrated *in vivo* and *in silico* approach

**DOI:** 10.1101/2025.07.24.666549

**Authors:** Md. Mohabbot Hossen, Sudip Bose, Rabindra Nath Acharyya, Apurba Kumar Barman, Sahria Rahman, Kingkar Prosad Gosh, Md. Arju Hossain, Shrabanti Dev, Asish Kumar Das

## Abstract

*Hylocereus polyrhizus,* locally known as red dragon fruit, is valued for its nutritional benefits including high levels of antioxidants and is gaining popularity as both a food and a medicinal plant. The present study addressed the *in vivo and in silico* chemopreventive potential of *Hylocereus polyrhizus* peel extract (HPPE) in DMBA-croton oil induced skin carcinogenic model mice. The mRNA expression level of pro-inflammatory cytokines and inflammatory mediators in tumor mass were estimated by real time-RT-qPCR. In addition, molecular docking and molecular dynamic simulation analyses were conducted on the reported compounds. In the *in vivo* chemopreventive activity assessment, the peel extract at 500 mg/kg was found effective in reducing total tumor number, yield, burden, incidence, and weight. Total proteins and endogenous antioxidants (GSH, SOD, CAT) levels in liver and skin tissues from mice were significantly (*P<0.05*) elevated. In addition, the HPPE at 500 mg/kg dose significantly reduced (*P<0.05*) the gene expression of pro-inflammatory cytokines such as TNF-*α*, IL-1*β*, IL-6, IL-18, and inflammatory mediators like TGF-*β*1, COX-2, and NFκ*Β*. In the molecular docking studies, reported compounds including Quercimeritrin, Rutin, and Kaempferol 3-O-β-D-glucopyranoside were identified as the top-performing compounds, with a docking score of - 7.4, -7.1 and -7.0 kcal/mol against TGF-*β*1 protein. This indicates stronger binding interactions compared to vincristine (-5.3 kcal/mol). In drug-likeness assessment, all compounds demonstrated the most favourable ADMET and pharmacokinetic profile. Furthermore, MDS data showed greater dynamic stability for Kaempferol 3-O-β-D-glucopyranoside and Rutin, while vincristine exhibited higher fluctuations. The results suggest that HPPE may serve as a potential inhibitor of skin carcinogenesis through upregulating endogenous antioxidants as well as suppressing different proinflammatory and inflammatory cytokines. Quercimeritrin, Rutin, and Kaempferol 3-O-β-D-glucopyranoside might be the probable leads responsible for this chemopreventive activity.

**Highlights:**

✓ *Hylocereus polyrhizus* peel extract (HPPE) demonstrated notable chemopreventive potential against skin cancer.
✓ HPPE significantly increased endogenous antioxidants (GSH, SOD, CAT) in liver and skin tissues, suggesting enhanced cellular defense mechanisms.
✓ Gene expression of pro-inflammatory cytokines (TNF-α, IL-1β, IL-6, IL-18) and inflammatory mediators (TGF-*β*1, COX-2, NFκB) were notably suppressed by HPPE.
✓ Quercimeritrin (CID: 5282160) demonstrated strong binding to TGF-*β*1 with a docking score of -7.4 kcal/mol, outperforming vincristine (-5.3 kcal/mol).
✓ Kaempferol 3-O-beta-D-glucopyranoside (CID: 5282102) and rutin (CID: 5280805) showed the best ADMET, pharmacokinetic profiles, and molecular stability, supporting their potential as lead compounds.

## Introduction

Cancer is one of the most common conditions of morbidity and mortality globally, and its different types present different clinical challenges. Skin cancer—melanoma, for instance— is well known for its aggressiveness and low survival rate [1]. Skin cancer is increasing day by day across the globe, and it’s a matter of great public health concern. In most parts of the world, as in the United States, skin cancer incidence continues to rise with an estimated 5.4 million new cases yearly [2]. Prognosis is still poor for patients with advanced disease, whereby distant metastases are met with a five-year survival of merely 16% and a median survival of four to six months [3].

In spite of developments in other modes of treatment like surgery, radiation, and chemotherapy, chemotherapy is still the most widespread method of treatment for skin cancers [4]. Its curative action, however, is typically negated by a wide array of side effects like myelotoxicity [5], cardiotoxicity [6], nephrotoxicity [7], pulmonary toxicity [8], immunosuppression, and alopecia [9]. The above shortcomings have generated scientific interest in alternative and complementary approaches, especially those of natural origin.

In the past several years, interest in the exploration of plant-based bioactive compounds with anticancer activity has grown [10]. Medicinal plants such as *Andrographis paniculata* and *Centella asiatica* have been reported to possess promising anticancer activities [11, 12]. Such activities have largely been credited to the action of phytoconstituents such as flavonoids and phenolics that are reported to disrupt carcinogen metabolism and enhance detoxification mechanisms [13, 14]. Exploiting such phytochemicals for chemoprevention is a new and promising approach in the fight against cancer [15].

One of those potential plants is *Hylocereus polyrhizus*, or red dragon fruit/pitaya. Belonging to the Cactaceae family, the fruit is originally from the American continent but is now being commercially grown in countries such as Bangladesh owing to its therapeutic and economic importance [16]. It has been regarded as a valuable medicinal plant in traditional Asian medicine and is widely utilized in treating as well as preventing a number of diseases [17].

Recent scientific research has supplied evidence of the biological activity of *H. polyrhizus* [18–21]. The peel of the fruit, routinely wasted, contains antioxidants like betacyanins, flavonoids, phenolics, and polyphenols [22]. Additionally, it contains essential vitamins (C, E, A), terpenoids, and bioactive compounds such as carotenoids and phytoalbumins which are responsible for its antioxidant and possible antibacterial activity. Different studies also described the fruit’s hypoglycemic, diuretic, anti-tumor, and antioxidative effects [24, 25], and its effectiveness in blood pressure, obesity, and dysentery [26].

Skin cancers are classified into melanoma due to dysregulation of melanocytes and non-melanoma skin cancers (NMSC), accounting for approximately 95% and are predominantly governed by genetic and environmental factors [27–30]. Inflammation is among the major biochemical pathways implicated in the development of melanoma and NMSC [31, 32]. Cytokines such as interleukins (ILs) and tumour necrosis factor (TNF) and anti-inflammatory markers such as TGF-β1 are the key players of inflammatory responses of the skin [33, 34]. NF-κB is a master regulator of inflammatory responses, controlling chemokine, cytokine, and adhesion molecule expression that takes part in cancer development [35].

While numerous studies on the phytochemical composition and in vitro bioactivity of *H. polyrhizus* have been carried out, detailed investigation into its anticancer potential— through in vivo and in silico approaches—has been limited. In silico approaches have transformed natural product research since these enable the prediction of bioactivity and speed up drug development with less cost and time [36].

In light of the growing need for safe and effective skin cancer therapeutics, the present study aims to evaluate the chemopreventive effects of *Hylocereus polyrhizus* peel extract (HPPE) using a DMBA/Croton oil-induced skin cancer mouse model. Both in vivo and in silico approaches were employed, with a particular focus on the modulation of mRNA expression of pro-inflammatory cytokines and inflammatory mediators.

## Materials and Methods

### Collection and extraction

Fruit peels from the *Hylocereus polyrhizus* were collected from the Jashore area in Bangladesh and confirmed by specialists at the Bangladesh National Herbarium in Dhaka (Accession no. DACB 85928). The collected peel was separated from the flesh and dried by shade drying. After grinding, the powder (800 g) was successively extracted with ethanol at room temperature, where the sample and solvent ratio was 1:5. A rotary evaporator was used to concentrate the extract, and the yield was 2.5%.

### Experimental animals and Ethical Statement

Young Swiss albino mice aged 4-5 weeks; weight 25-30 g was used for the experiment. The mice were procured from Pharmacy Department, Jahangirnagar University, Bangladesh. The animals were kept under ideal environmental conditions in the animal house of the Pharmacy Discipline, Khulna University. We strictly adhered to the Ethical Committee’s guidelines (approval number: KUAEC-2022/03/02) for the handling and care of the experimental mice.

### Chemicals

We bought 7,12-dimethylbenz (α) anthracene (DMBA), Croton oil, Tris-HCl, BSA (Bovine Serum Albumin), reduced glutathione (GSH), TCA (trichloroacetic acid), and DTNB (5,5 dithiobis (2-nitrobenzoic acid)) from Sigma Chemicals Co. in St. Louis, MO, USA. Vincristine was obtained from Beacon Pharmaceuticals Limited in Mymensingh, Bangladesh. We purchased the PureLink® RNA Mini Kit, RevertAid First Strand cDNA Synthesis Kit, and SYBR Green master mix from Thermo Fisher Scientific (USA). Forward and reverse primers for GAPDH, TNF-α, NF-κB, IL-1β, IL-6, IL-18, and TGF-*β*1 were purchased from Invent Technologies Limited, Dhaka, Bangladesh. All other chemicals were used in analytical grade.

### Experimental design

#### Evaluation of chemopreventive potential

The chemopreventive effect of HPPE was checked by chemically induced skin carcinogenesis in Swiss albino mice as described by Elguindy [37] with minor modification. Sixty experimental mice were chosen randomly before the experiment, and the back areas of the mice’s (3x3 cm²) skin were shaved. The experimental mice were divided into five groups: control group, carcinogenic control group, standard group, and two test groups that received HPPE at doses of 250 mg/kg and 500 mg/kg body weight, with 15 mice in each HPPE group and ten mice in the other groups. The control group received 10 ml/kg of 2% Tween-80 in water orally and 50 µL of acetone topically applied to their shaved backs. The standard and carcinogenic control groups received a single topical application of DMBA (100 µg in 50 µL of acetone per animal) on their shaven back. Two weeks afterwards, 0.1 ml of 1% croton oil in acetone was topically applied on each alternative day until the end of the experiment at 16 weeks. The standard group were given a dose of 500 µg/kg vincristine once a week for last one month. We administered 250 and 500 mg/kg HPPE extract orally to two test groups before 7 days of DMBA-induced carcinogenesis, and continued this daily until the experiment terminated.

Throughout the investigation, we closely observed the mice and recorded their body weight weekly basis. We recorded the papilloma appearance rates in each mouse across all groups on a daily basis. The ‘Tumour Incidence’ (the number of mice with at least one tumour), ‘Tumour Yield’ (the average number of papillomas per mouse), and ‘Tumour Burden’ (the average number of tumours per tumour-bearing mouse) were noted carefully.

#### Biochemical study

We followed Sharma’s previously described method with minor modification [33] to prepare the tissue homogenate. After sacrificing the animals, the skin was taken from the chosen dorsal area and carefully cleaned with cooled 0.9% NaCl (pH 7.4) solution. Following the weighing of the tissue (skin), a 10% tissue homogenate was prepared using a portion of the sample and 0.15 M Tris-HCl (pH 7.4) by a homogenizer. We then centrifuged the homogenate at 2500 rpm for 10 minutes. The resulting supernatant was used to calculate the concentration of total protein, reduced glutathione (GSH), SOD and catalase.

#### Estimation of total protein and endogenous antioxidants (GSH, SOD, catalase) concentration

Estimation of total protein: As a standard, bovine serum albumin was used to conduct the experiment [38]. Briefly, a 1.0 mL sample was diluted with 0.9 mL reagent A (2 g sodium potassium tartrate and 100 g sodium carbonate in 500 mL 1N NaOH) and cooled down to room temperature after being incubated for 10 minutes at 50°C. Reagent B of 0.1 mL (2 g sodium potassium tartrate, 1 g copper sulphate in 90 mL H₂O and 10 mL 1N NaOH) was then added to it and incubated for 10 min at room temperature. Finally, 3 mL reagent C (1 volume Folin-Ciocalteau reagent diluted with 15 volumes of water) was added, mixed and incubated for 10 min in the 50°C water bath. In order to measure the absorbance, a UV-VIS spectrophotometer was used at 650 nm.

GSH estimation: GSH level was estimated following the method described by Alejandro et al. [39]. Initially, 0.1 ml of tissue homogenate was precipitated with 100 µL of 5% TCA (trichloroacetic acid). For complete protein precipitation, the mixture was thoroughly mixed before centrifuging. After that, 0.1 ml of supernatant, 2.0 ml of 0.6 mM DTNB [5, 5 dithiobis (2-nitrobenzoic acid)] reagent and 0.2 M phosphate buffer (pH 8.0) were added to make up to a final volume of 4.0 ml. At 412 nm, the absorbance was measured against a blank.

SOD Estimation: SOD activity was measured following the procedure as described by Alejandro et al. [39]. In brief, 2.65 mL of carbonate buffer was taken, and 300 µL of tissue homogenate was mixed with it. After that, 0.02 ml EDTA was added, followed by 0.03 ml of epinephrine. Finally, absorbance was taken at 480 nm.

Catalase estimation: The CAT activity was measured following the method described by Aebi et al. [40]. First, 2.8 ml 30 mM H_2_O_2_ was placed in a blind tube and 0.2 ml phosphate buffer was added to it. Then 2.8 ml 30 mM H_2_O_2_ was added to the sample tube. To both of these tubes, 0.2 ml enzyme was added, and the tubes were mixed well through vortexing. The absorbances at 240 nm were read twice at 30 s intervals to determine the activity.

#### Study of different serum biochemical parameters

Various biochemical parameters, i.e., Serum Glutamic Pyruvic Transaminase (SGPT), serum Glutamic-Oxaloacetic Transaminase (SGOT), urea, bilirubin, creatinine, total cholesterol, High-Density Lipoprotein (HDL), Low-Density Lipoprotein (LDL), and Triglycerides (TG) were assessed following the established protocol. At the end of experiment, the animals were anaesthetized, and their blood samples were collected through Jugular vein without anticoagulant (EDTA). For biochemical analysis, blood without additive was centrifuged at 1500 rpm for 20 min, and serum was separated.

#### Measurement of mRNA Expression induce by DMBA-croton oil

RNA was extracted from skin tissue samples of five different groups (negative control, carcinogenic control, standard, test 1, 250 mg/kg, and test 2, 500 mg/kg) using the PureLink® RNA Mini Kit from Thermo Fisher Scientific, USA. After that, first strand cDNA was made using the RevertAid First Strand cDNA Synthesis Kit from Thermo Fisher Scientific, USA following manufacturer’s protocol.

#### RT-qPCR

RT-qPCR was conducted with PowerUp™ SYBR™ Green Master Mix (Thermo Fisher Scientific, USA). The primers used in the experiment are shown in Table 1. Glyceraldehyde-3-phosphate dehydrogenase (GAPDH) was used as the internal control. The 2-ΔΔCT method was used to assess the fold changes in messenger RNA (mRNA) levels in all groups.

**Table 1:**
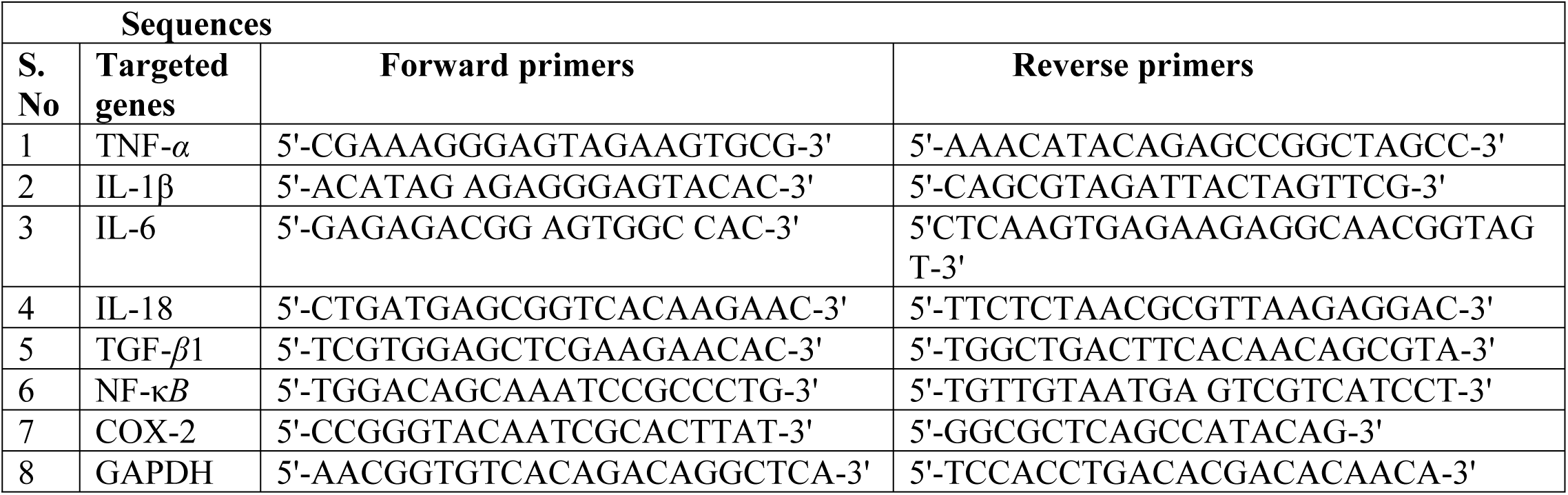
Primers used in real time-RT-qPCR technique.

#### Statistical analysis

Results were expressed as the mean ± standard error of means (SEM). Dunnett’s t test was used to determine statistical significance. Post-hoc Tukey tests were used to compare two treatment groups side by side. *P<0.05* was considered statistically significant.

#### In silico approaches

RT-qPCR analysis showed significant upregulation of inflammatory genes including TNF-α, IL-1β, IL-6, IL-18, NF-κ*B*, COX-2, and notably TGF-β1, which was the most elevated.

Therefore, molecular docking focused on TGF-β1 due to its key role in skin carcinogenesis and the availability of a suitable crystal structure for structure-based drug design [41].

#### Molecular docking

We chose ten (10) potential drug molecules from previous literature [42]. The 3D shapes of these compounds were downloaded from the PubChem database (https://pubchem.ncbi.nlm.nih.gov/) and prepared for docking using AutoDock Tools 1.5.6 [43]. The energy minimisation of the ligands was carried out using Avogadro software with the MMFF94 force field and steepest descent algorithm to ensure a stable conformation [44].

The 3D structure of the TGF-*β*1 protein (PDB ID: 6B8Y) was obtained from the RCSB Protein Data Bank server (https://www.rcsb.org/). The receptor proteins were preprocessed using AutoDock Vina by adding charges and reducing energy. Using AutoDock Tools 1.5.6, heteroatoms, water molecules, and bound ligands were removed from the protein structure. Polar hydrogens were added to the protein for docking simulations. A grid box was made around the TGF-*β1* protein’s (PDB ID: 6B8Y) binding site to set the area where ligands can dock. The grid box size (126x126x126) and the grid spacing (0.375 Å) were selected based on the size of the binding site and the required resolution. The centre of the grid box (x: -2.227; y: 95.968 and z: 114.882) was positioned at the coordinates corresponding to the active site of the protein.

We performed molecular docking using AutoDock Vina, utilizing the previously prepared ligands and the receptor protein. The exhaustiveness parameter was set to 50 to balance search thoroughness and computational efficiency. Docking positions were created for each ligand, and the strength of their connection to the protein (measured in kcal/mol) was calculated for the protein-ligand pairs. Interaction analysis was conducted using the visualisation tool, Biovia Discovery Studio Visualiser v21, to examine non-covalent interactions.

#### ADMET and drug-likeness properties analysis

The pharmacokinetic properties of the highest docking scored candidates were evaluated through comprehensive ADMET (Absorption, Distribution, Metabolism, Excretion, and Toxicity) analysis using their optimised molecular structures in SMILES format. The ADMET properties were predicted using online databases such as SwissADME (https://www.swissadme.ch/) and pkCSM (https://biosig.lab.uq.edu.au/pkcsm/) servers. The evaluation included human intestinal absorption (HIA), with compounds showing an HIA ≥ 30% considered suitable for effective absorption. The ability of compounds to cross the blood-brain barrier was checked using LogBB values, where those with a LogBB of 0.3 or higher were seen as possibly able to pass through the barrier. We also subjected the compounds to Lipinski’s Rule of Five analysis to evaluate their drug-likeness. The criteria included having fewer than 10 hydrogen bond acceptors, fewer than 5 hydrogen bond donors, a molecular weight of less than 500 Da, and a LogP value of less than 5 [45].

#### Molecular dynamics simulation (MDS) studies

The selected ligand molecules’ conformational behavior and protein stability were examined using a dynamics simulation. We chose Quercimeritrin (CID: 5282160), Rutin (CID: 5280805), and Kaempferol 3-O-β-D-glucopyranoside (CID: 5282102), and for MDS after investigating binding affinity, drug-likeness, and pharmacokinetic features. The "Desmond v3.6 Programme" (https://www.schrodinger.com/) was used to execute a 200 ns MDS (paid version) in a Linux environment to ascertain the binding consistency of the protein-ligand complex structures [46]. Using this setup, the shape of the orthorhombic periodic bounding box is divided with a 10 Å space for the water boundary in a set volume using the TIP3P water model. To balance the electric charge in its structure, sodium (Na⁺) and chloride (Cl⁻) ions (0+ and 0.15 M salt) were selected and randomly spread throughout the surrounding water [46, 47].

After creating solvated protein structures with different agonist combinations, the system’s setup was simplified, and it effectively followed the procedure using the OPLS3e force field constants included in the Desmond package. A steady atmospheric pressure of 1.01325 bar and a temperature of 300 K were maintained for each NPT assembly, which used a uniform method and included 50 pauses of 50 picoseconds with an efficiency of 1.2 kcal/mol. The accuracy of the MD simulation was checked during the whole process using the Simulations Interaction Diagram (SID) from the Desmond tools in the Schrödinger suite [1]. By using data from root mean square deviation (RMSD), root mean square fluctuation (RMSF), radius of gyration (Rg), solvent accessible surface area (SASA), intramolecular bond analysis, polar surface area (PSA), and molecular surface area (MolSA), it was possible to assess the relative solidity of specific protein-ligand interaction combinations [1].

#### MM-GBSA binding free energy analysis

The MM-GBSA (Molecular Mechanics Generalized Born Surface Area) method was employed using GROMACS to estimate the binding free energies (ΔG) of the ligand-receptor complexes. Followed by molecular dynamics (MD) simulations in GROMACS, about 8-10 Å explicit water boundaries were generated for system equilibration. Snapshots from the equilibrated MD phase were used for MM-GBSA calculations, where ΔG was computed as the sum of molecular mechanics energy (ΔE_MM) and solvation free energy (ΔG_solvation), with the entropy term (TΔS) not explicitly calculated [1]. Binding affinities were averaged across multiple snapshots, with negative ΔG values indicating thermodynamically favorable interactions and highlighting the binding potential of the tested compounds.

## Results

### Chemopreventive activity

The results showed that the total number of tumors, their size, and weight were higher in the carcinogenic control group, highlighting the skin cancer model caused by DMBA-croton oil. All treatment groups showed a decrease in these values (Table 2). In comparison to the carcinogenic group, the higher dose of HPPE extract (500 mg/kg) significantly reduced the total number of tumors by 1.5 times and the total tumor weight by 1.4 times. Additionally, it lowered the overall tumor burden and occurrence, demonstrating its effectiveness in preventing skin cancer.

**Table 2:** Effect of HPPE on different tumor parameters.

| Parameters | Control | Carcinogenic Control | Standard | HPPE (250 mg/kg) | HPPE (500 mg/kg) |
| --- | --- | --- | --- | --- | --- |
| Total tumor no. | 0 | 48 | 23 | 39 | 29 |
| Total tumor weight (mg) | 0 | 1680 | 665 | 1505 | 1155 |
| Tumor yield | 0 | 8 | 2.09 | 4.87 | 2.9 |
| Tumor burden | 0 | 8 | 2.55 | 4.87 | 3.62 |
| Tumor incidence | 0% | 100% | 82% | 100% | 90% |

### Estimation of total protein and endogenous antioxidants (GSH, SOD, catalase) concentration

Total protein and different endogenous antioxidant molecule (GSH, SOD, and CAT) concentrations in mice tumor tissue and liver were assessed. We observed that mice treated with DMBA/croton oil had remarkably decreased levels of total protein and endogenous antioxidant molecules compared to the control group (Table 3). In contrast, the treatment groups receiving HPPE (both 250 mg/kg and 500 mg/kg) showed an increase in total protein and endogenous antioxidant molecules compared to the carcinogenic control showing that it has a beneficial effect on endogenous antioxidant.

**Table 3:** Effect of HPPE on total protein, GSH, SOD and CAT concentration.

| Parameters | Sample | Control | Carcinogenic Control | Standard | HPPE (250 mg/kg) | HPPE (500 mg/kg) |
| --- | --- | --- | --- | --- | --- | --- |
| **Total protein (µg/mg tissue)** | Tumor tissue | 19.35 ± 0.58 | 13.76 ± 0.27<sup>#</sup> | 17.83 ± 0.77 *** | 14.63 ± 0.45 | 15.92 ± 0.26*** |
|  | Liver | 21.96 ± 0.30 | 14.65 ± 0.11<sup>#</sup> | 21.39 ± 0.73*** | 14.74 ± 0.32 | 16.83 ± 0.58** |
| **GSH (µg/mg protein)** | Tumor tissue | 964.26 ± 22.31 | 682.25 ± 58.4<sup>#</sup> | 876.57 ± 41.9** | 780.00 ± 46.78 | 859.86 ± 57.8* |
|  | Liver | 985.77 ± 16.29 | 731.97 ± 16.7<sup>#</sup> | 873.39 ± 11.5 *** | 772.53 ± 5.50 ** | 795.11 ± 8.11** |
| **SOD (U/mg protein)** | Tumor tissue | 85.49 ± 2.53 | 27.41 ± 3.16<sup>#</sup> | 73.11 ± 1.75 *** | 33.87 ± 1.54 | 55.84 ± 0.45*** |
|  | Liver | 75.22 ± 1.09 | 28.75 ± 4.25<sup>#</sup> | 67.93 ± 3.02 *** | 39.95 ± 2.09 | 62.03 ± 3.03*** |
| **CAT (U/mg protein)** | Tumor tissue | 226.79 ± 1.02 | 152.52 ± 1.69<sup>#</sup> | 221.67 ± 3.81 *** | 171.14 ± 6.57 ** | 180.62 ± 4.14*** |
|  | Liver | 224.52 ± 3.15 | 155.92 ± 8.3<sup>#</sup> | 209.17 ± 2.41 *** | 158.16 ± 2.86 | 186.27 ± 2.37 ** |

Values are expressed as mean ± standard error of mean; <sup>#</sup> $P \leq 0.05$ vs control; \* $P \leq 0.05$ vs carcinogenic control; \*\* $P \leq 0.01$ vs carcinogenic control, \*\*\* $P \leq 0.001$ vs carcinogenic control.

### Serum biochemical parameters

In this experiment, the carcinogenic control group exhibited significantly higher levels of SGPT, SGOT, urea, bilirubin, creatinine, and LDL, along with lower HDL levels, compared to the control group. The standard drug significantly reduced these elevated biochemical parameters (Table 4). The HPPE treatments, particularly at 500 mg/kg, also showed a reduction in these elevated parameters, indicating their potential to mitigate the biochemical alterations induced by the carcinogenic condition (Table 4).

**Table 4:**
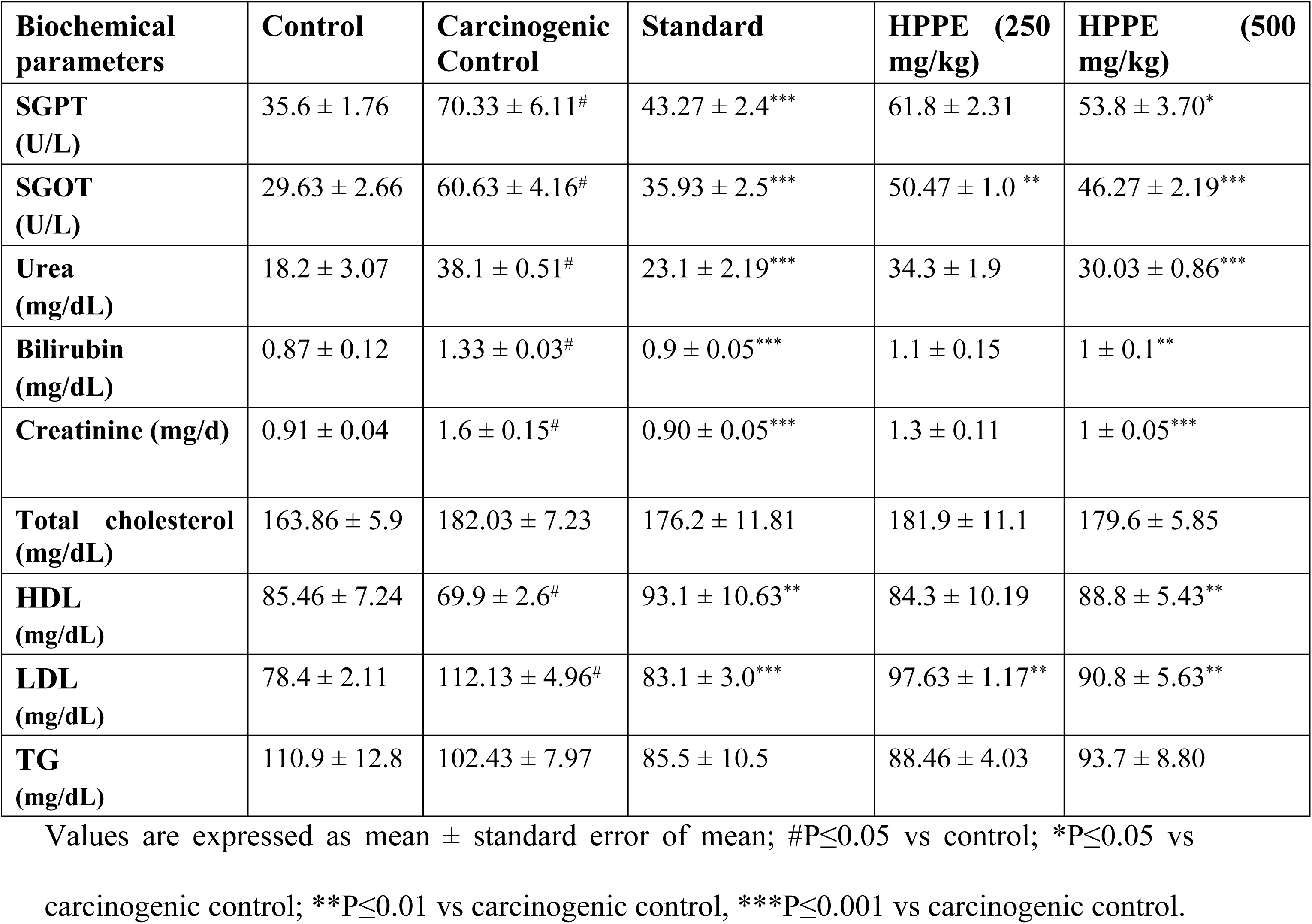
Effect of HPPE on different serum biochemical parameters.

| Biochemical parameters | Control | Carcinogenic Control | Standard | HPPE (250 mg/kg) | HPPE (500 mg/kg) |
| --- | --- | --- | --- | --- | --- |
| SGPT (U/L) | 35.6 ± 1.76 | 70.33 ± 6.11<sup>#</sup> | 43.27 ± 2.4<sup>***</sup> | 61.8 ± 2.31 | 53.8 ± 3.70<sup>*</sup> |
| SGOT (U/L) | 29.63 ± 2.66 | 60.63 ± 4.16<sup>#</sup> | 35.93 ± 2.5<sup>***</sup> | 50.47 ± 1.0<sup>**</sup> | 46.27 ± 2.19<sup>***</sup> |
| Urea (mg/dL) | 18.2 ± 3.07 | 38.1 ± 0.51<sup>#</sup> | 23.1 ± 2.19<sup>***</sup> | 34.3 ± 1.9 | 30.03 ± 0.86<sup>***</sup> |
| Bilirubin (mg/dL) | 0.87 ± 0.12 | 1.33 ± 0.03<sup>#</sup> | 0.9 ± 0.05<sup>***</sup> | 1.1 ± 0.15 | 1 ± 0.1<sup>**</sup> |
| Creatinine (mg/d) | 0.91 ± 0.04 | 1.6 ± 0.15<sup>#</sup> | 0.90 ± 0.05<sup>***</sup> | 1.3 ± 0.11 | 1 ± 0.05<sup>***</sup> |
| Total cholesterol (mg/dL) | 163.86 ± 5.9 | 182.03 ± 7.23 | 176.2 ± 11.81 | 181.9 ± 11.1 | 179.6 ± 5.85 |
| HDL (mg/dL) | 85.46 ± 7.24 | 69.9 ± 2.6<sup>#</sup> | 93.1 ± 10.63<sup>**</sup> | 84.3 ± 10.19 | 88.8 ± 5.43<sup>**</sup> |
| LDL (mg/dL) | 78.4 ± 2.11 | 112.13 ± 4.96<sup>#</sup> | 83.1 ± 3.0<sup>***</sup> | 97.63 ± 1.17<sup>**</sup> | 90.8 ± 5.63<sup>**</sup> |
| TG (mg/dL) | 110.9 ± 12.8 | 102.43 ± 7.97 | 85.5 ± 10.5 | 88.46 ± 4.03 | 93.7 ± 8.80 |

Values are expressed as mean ± standard error of mean; <sup>#</sup>P≤0.05 vs control; <sup>\*</sup>P≤0.05 vs carcinogenic control; <sup>\*\*</sup>P≤0.01 vs carcinogenic control, <sup>\*\*\*</sup>P≤0.001 vs carcinogenic control.

### mRNA Expression

We assessed the mRNA expression levels of various pro-inflammatory cytokines, including TNF-*α*, IL-1*β*, IL-6, and IL-18, as well as inflammatory mediators like TGF-*β*1, COX-2, and NF-κB, using Real Time RT-PCR. Our data exhibited that the mRNA expression of both pro-inflammatory cytokines and inflammatory mediators was expressed more than 4.5 times in the carcinogenic group compared to normal mice (control group), as shown in Fig. 1. The standard drug vincristine sulfate significantly lowered the mRNA expression of targeted pro-inflammatory cytokines and inflammatory mediators compared to the carcinogenic group. Importantly, a significant reduction in the expression of pro-inflammatory cytokines and inflammatory mediators was observed with a high dose of HPPE (500 mg/kg), as compared to untreated carcinogenic mice, as shown in Fig. 1.

**Fig. 1:**
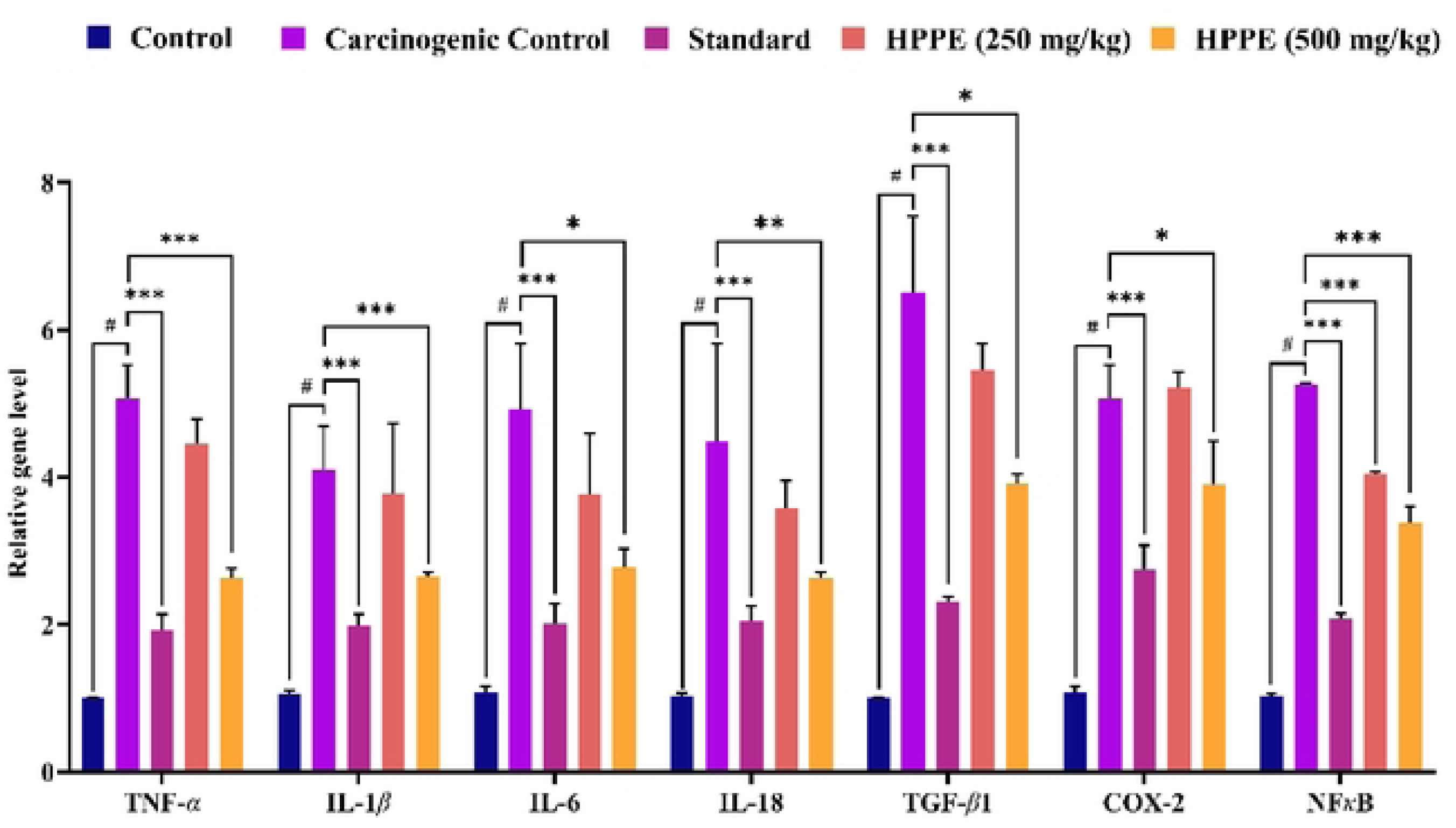
Effect of *Hylocereus polyrhizus* peels on mRNA expression level of the pro­ inflammatory cytokines (TNF-a, IL-lP, IL-6 and IL-18) and the inflammatory mediators (TGF-/Jl, COX-2 & NFKB) in DMBA induced skin carcinogenic mice. Values are expressed as mean ± standard error of mean, #P<0.05 vs control; *P<0.05 vs carcinogenic control; **P<0.01 vs carcinogenic control, ***P<0.001 vs carcinogenic control.

### In silico Analysis

#### Molecular Docking and Binding Affinity Analysis

A molecular docking study targeting TGF-β1 (PDB ID: 6B8Y) revealed that CID: 5282160 (Quercimeritrin) exhibited the highest binding affinity with a docking score of -7.4 kcal/mol, followed by CID: 5282102 (Kaempferol 3-O-β-D-glucopyranoside) and CID: 5280805 (Rutin) with scores of -7.1 and -7.0 kcal/mol, respectively. The control compound, vincristine, had a docking score of -5.3 kcal/mol (Supplementary Table S1). These results indicate that Quercimeritrin, Kaempferol 3-O-β-D-glucopyranoside, and Rutin could be effective options for blocking TGF-*β*1 to help treat skin cancer caused by DMBA-carton oil.

The molecular protein-ligand interaction analysis highlights significant differences in binding interactions between the control and reported phytochemicals (Supplementary Table S2). In Rutin (Fig. 2A & 3A), the ligand demonstrates multiple hydrogen bonds with critical residues such as Ser-148, Glu-127, and Asp-146, which are crucial for stabilizing the ligand in the binding pocket. Additionally, carbon-hydrogen bonds and polar interactions enhance overall binding strength, with bond distances ranging from 2.2 Å to 4.5 Å, indicating strong anchoring. Kaempferol 3-O-β-D-glucopyranoside (Fig. 2B & 3B) has important hydrogen bonds with Lys-79, Ile-80, and Leu-135, along with strong hydrophobic and π-sigma interactions, which suggest it goes deeper into the protein’s hydrophobic areas. Besides, Quercimeritrin (Fig. 2C and 3C) showcases hydrogen bonding with Ser-148, Asn-116, and Asp-146 while π-stacking interactions with Phe-127, which collectively enhance the ligand’s binding stability through aromatic and polar interactions. In comparison, the control ligand vincristine (Fig. 2D) mainly connects with Asp-146 and Arg-149 using a few hydrogen bonds and weaker π-alkyl interactions, with bond distances reaching up to 4.8 Å, showing that its binding is relatively weaker.

**Fig. 2:**
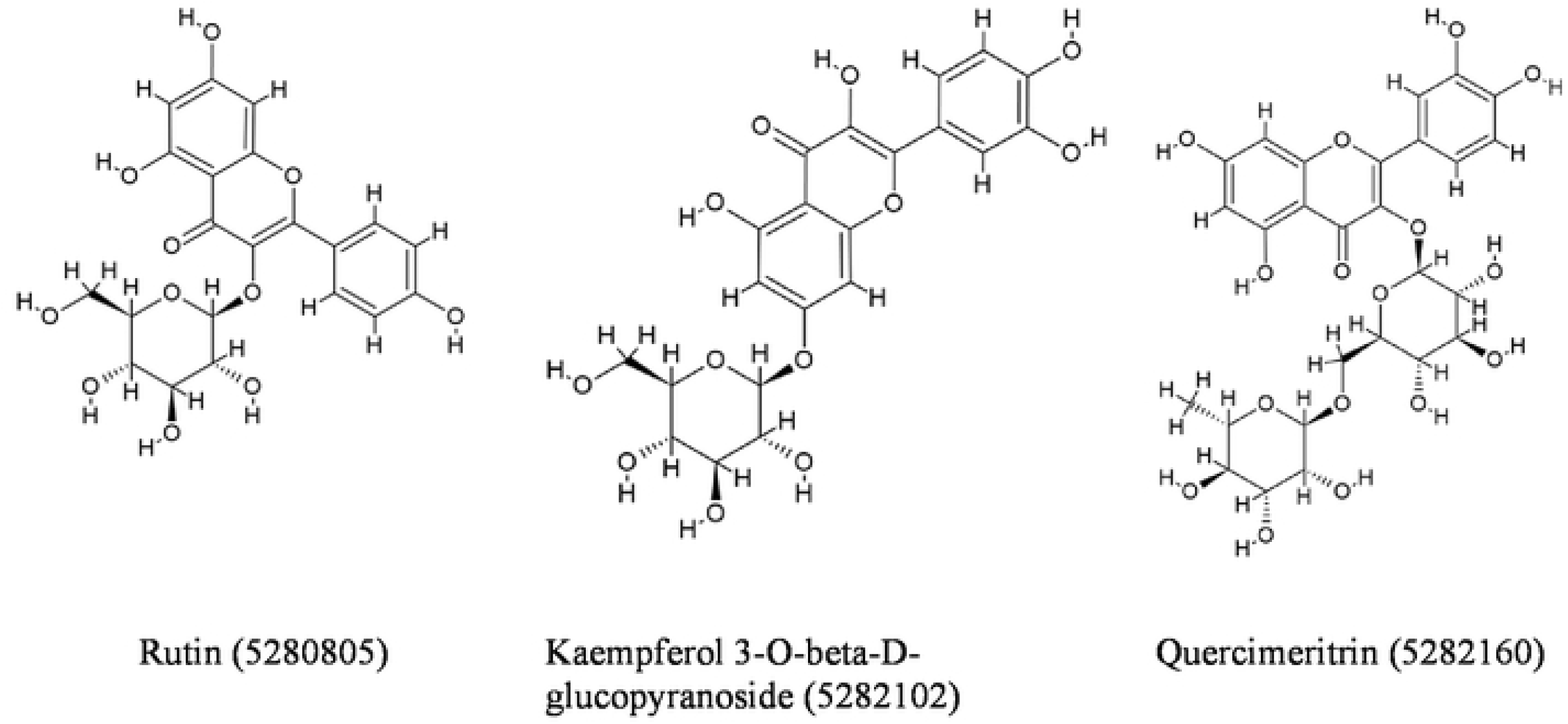
Structure of identified compounds that exhibited highest binding score against TGF-Pl (PDB ID: 6B8Y) protein (A) CID: 5280805, (B) CID: 5282102 & (C) CID: 5282160.

**Fig. 3:**
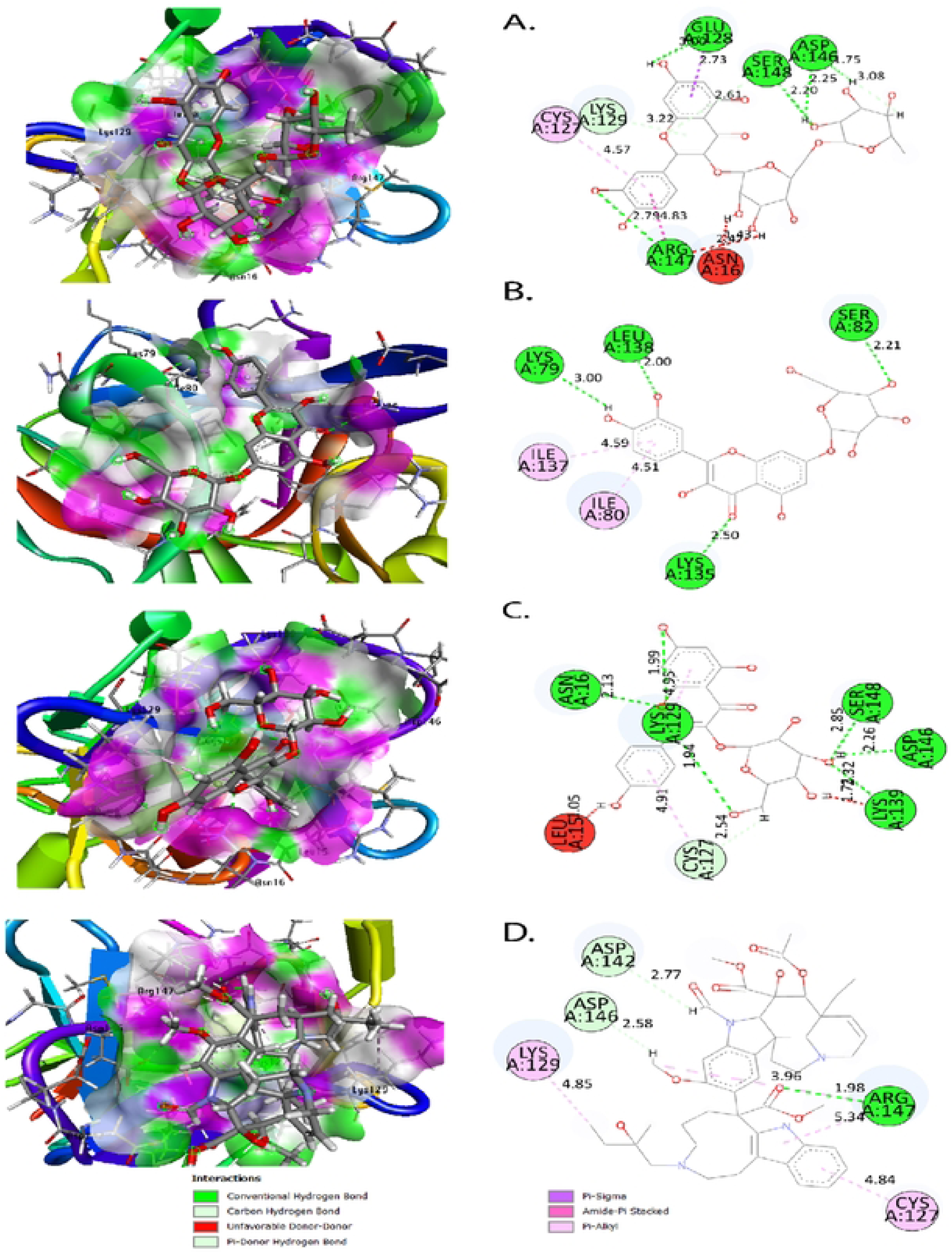
3D and 2D molecular binding interaction ofTGF-Pl (PDB ID: 6B8Y) against (A) phytocompound CID: 5280805, (B) phytocompound CID: 5282102, (C) phytocompound CID: 5282160, and (D) control ligand Vincristine.

#### ADMET and drug-likeness properties analysis

The ADMET properties of the tested compounds such as Quercimeritrin, Kaempferol 3-O-β-D-glucopyranoside, and Rutin were assessed for their drug-likeness and potential pharmacokinetic behaviour (Table 5). All compounds exhibited limited skin permeability (-2.735) and moderate Caco-2 permeability values. Volume of distribution (Vd) values indicated moderate tissue distribution, while blood-brain barrier (BBB) and central nervous system (CNS) permeability scores were negative across all compounds, suggesting poor central nervous system penetration. Metabolic stability was confirmed, as none of the compounds acted as substrates or inhibitors of CYP2D6 or CYP3A4 enzymes. Total clearance was positive for all compounds except Rutin. Toxicity predictions showed no AMES toxicity, hepatotoxicity, or skin sensitization risks. Physicochemical evaluations revealed molecular weights ranging from 448.38 to 770.69 Da, with hydrogen bond acceptor and donor counts within moderate ranges. The solubility of all compounds was categorized as "soluble," and lipophilicity values (Log Po/w) were low or negative. Despite some Lipinski rule violations, all compounds had a bioavailability score of 0.17 (Table 5).

**Table 5:**
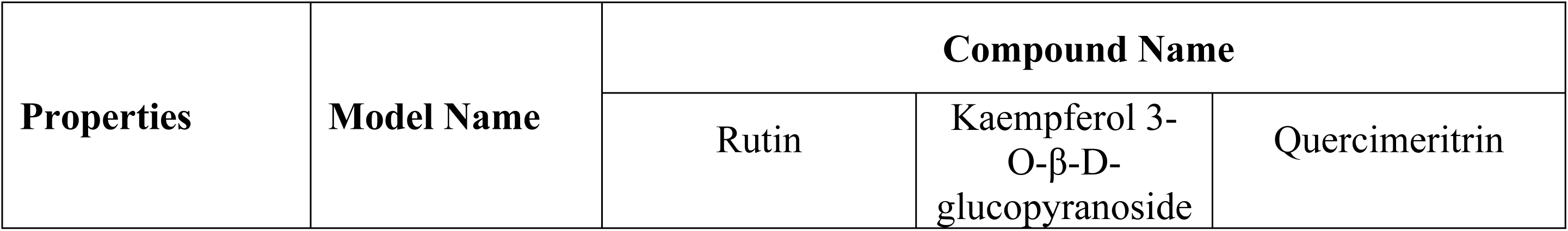
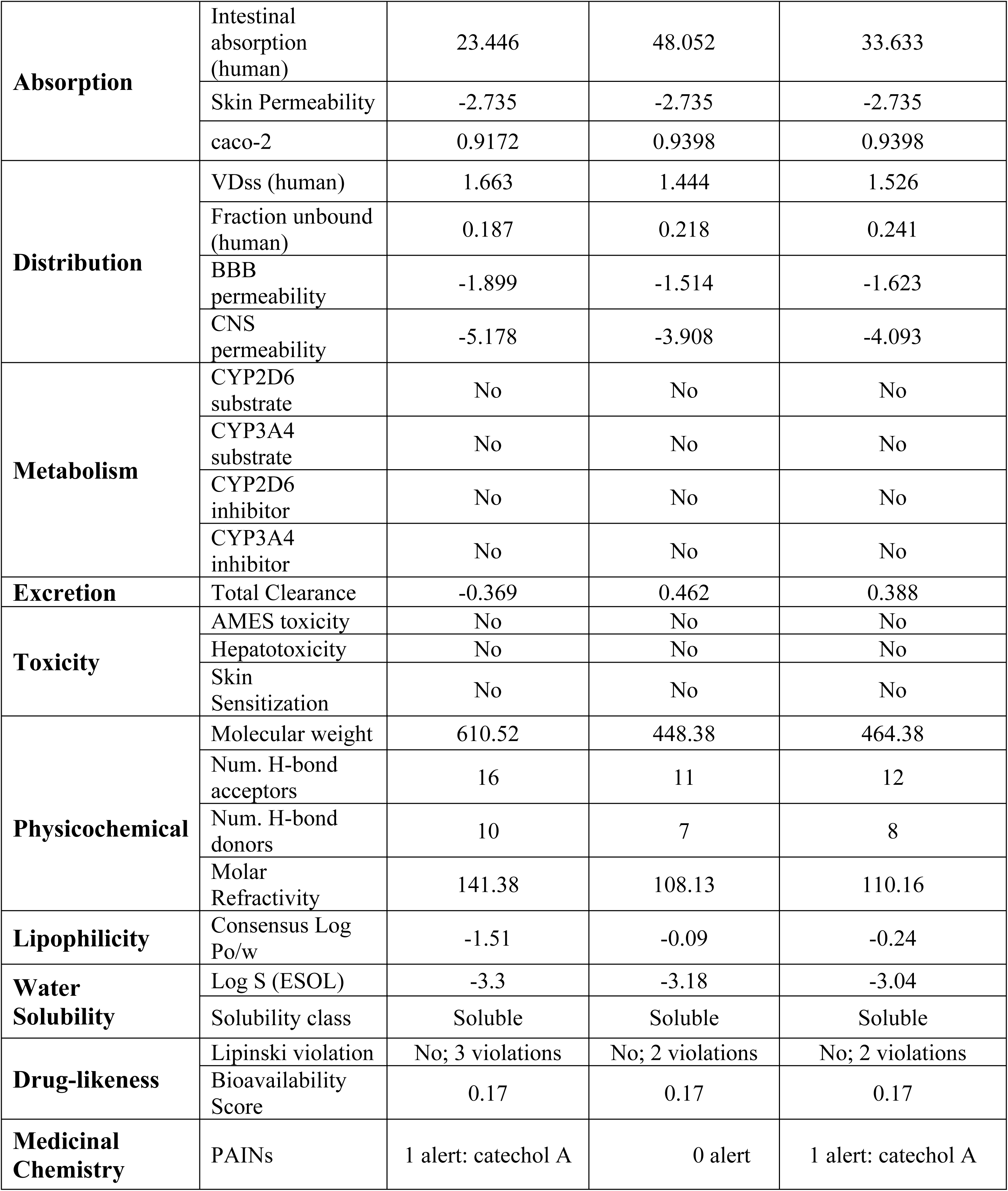
ADMET, Physiochemical, Lipophilicity and Drug-likeness properties analysis of reported compounds.

### Molecular dynamics simulation (MDS) studies

#### Root Mean Square Deviation (RMSD) and Fluctuation Analysis

The RMSD analysis (Fig. 4A) highlights the dynamic stability of the protein-ligand complexes over a 200 ns simulation. The control ligand (vincristine) displays the highest fluctuation, with RMSD values peaking around 3.5 Å, indicating a less stable binding interaction with the target protein. In contrast, the complexes with CID: 5280805 (Rutin) and CID: 5282102 (Kaempferol 3-O-β-D-glucopyranoside) demonstrate more stable behaviour, stabilizing around 2.5 Å after 50 ns, suggesting strong ligand-protein interactions and consistent conformations. In addition, CID: 5282160 (Quercimeritrin) shows moderate fluctuations between 2.5 Å and 3.0 Å, indicating intermediate stability compared to the control and the other two experimental ligands. These results imply that CID: 5280805 (Rutin) and CID: 5282102 (Kaempferol 3-O-β-D-glucopyranoside) form more stable complexes than the vincristine drug.

**Fig. 4:**
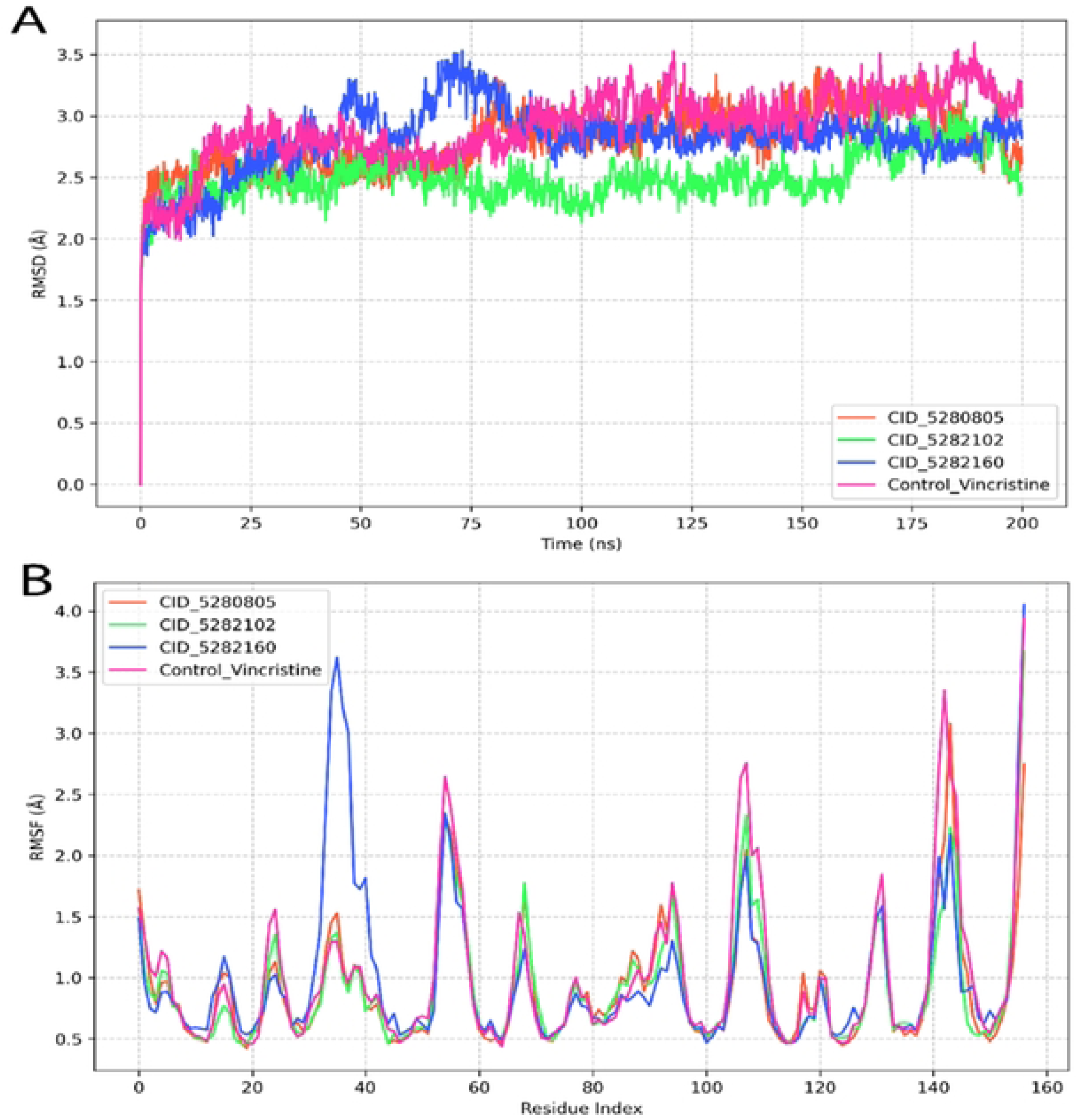
(A) Root Mean Square Deviation (RMSD) plot showing the structural stability of the protein backbone in complex with different ligands (CID_5280805, CID_5282102, CID_5282160) and a control (Vincristine) over a 200 ns simulation period. (B) Root Mean Square Fluctuation (RMSF) plot illustrating residue-wise flexibility of the protein. The highest fluctuations are observed in the loop regions, particularly around residues 40, 100, and 50.

The RMSF analysis (Fig. 4B) provides a residue-level view of the fluctuations across the protein. The control complex (vincristine) shows significant fluctuations, particularly in regions between residue indices 40–50 and 140–160, suggesting increased conformational flexibility. In comparison, CID: 5280805 (Rutin) and CID: 5282102 ((Kaempferol 3-O-β-D-glucopyranoside) exhibit minimal fluctuations across most residues, indicating tighter binding and reduced flexibility, which correlates with their lower RMSD values. In addition, CID: 5282160 shows moderate fluctuations, particularly at the C-terminal region, reflecting its slightly less stable binding. In a nutshell, the reduced flexibility and lower RMSD values observed for CID: 5280805 (Rutin) and CID: 5282102 (Kaempferol 3-O-β-D-glucopyranoside) suggest that these ligands may have higher binding affinity and stability compared to the control.

#### Radius of gyration (Rg) and solvent accessible surface area (SASA) analysis

The Rg was tracked for all ligand-protein complexes during a 200 ns molecular dynamics simulation (Fig. 5A). The control (vincristine) exhibited relatively higher fluctuations throughout the simulation, with an average Rg of approximately 15.4 Å. The CID: 5280805 (Rutin) and CID: 5282160 (Quercimeritrin) complexes showed the most consistent behavior, keeping Rg values below 15.35 Å, which means they are more compact and stable in their structure. CID: 5282102 (Kaempferol 3-O-β-D-glucopyranoside) showed slightly higher Rg fluctuations, suggesting intermediate stability. These results suggest that CID: 5280805 (Rutin) and CID: 5282160 (Quercimeritrin) maintain structural compactness better than the control under dynamic conditions.

**Fig. 5:**
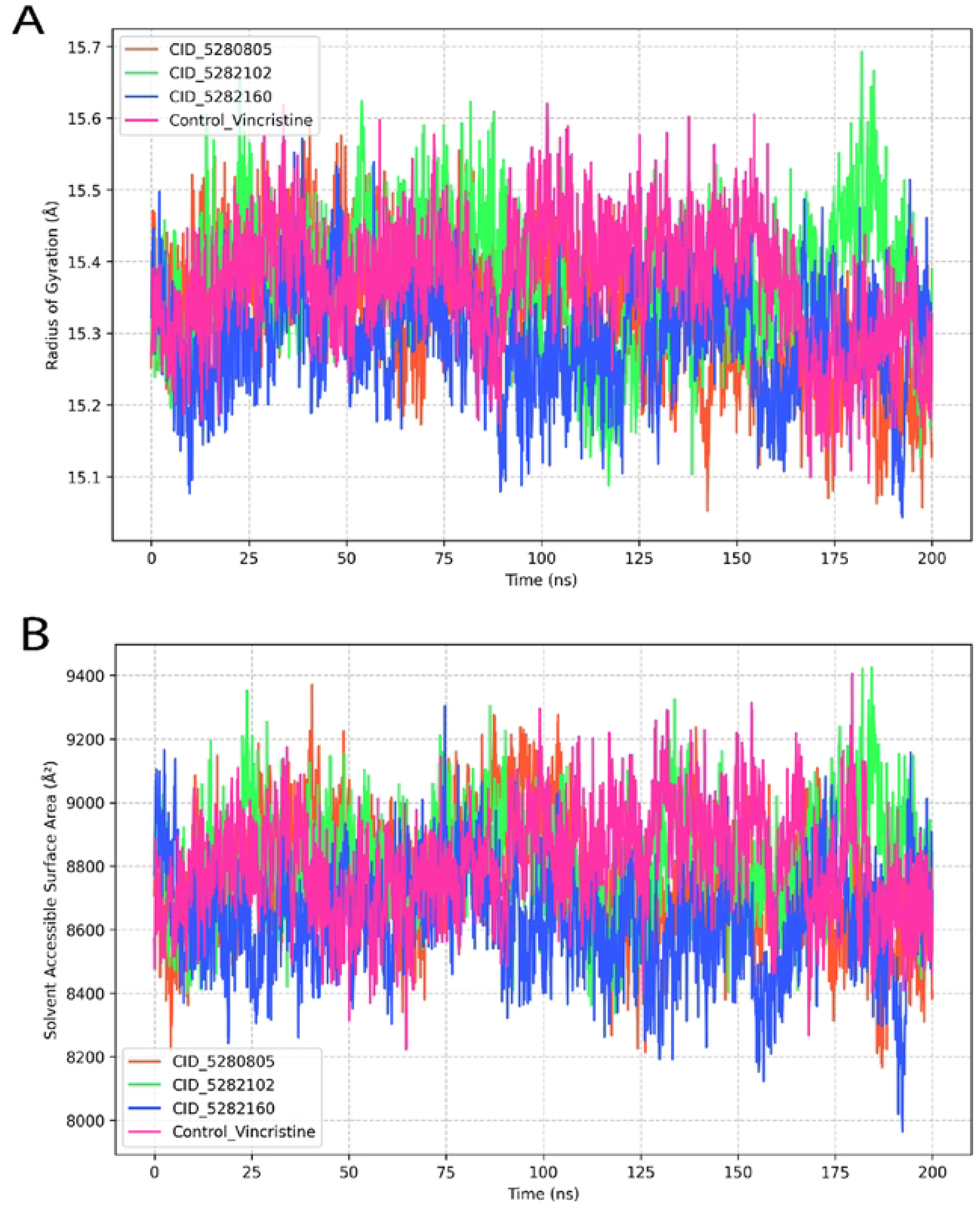
(A) Radius of gyration (Rg) plots showing compactness of protein-ligand complexes over 200 ns; all systems remain relatively stable with minor fluctuations. (B) Solvent accessible surface area (SASA) plots indicating dynamic surface exposure; lower SASA value suggesting tighter packing.

The SASA values for all complexes were evaluated to assess changes in solvent exposure (Fig. 5B). CID: 5282160 (Quercimeritrin) displayed the lowest SASA values, averaging around 8,200 Å², indicating reduced solvent exposure. In contrast, the control and CID: 5282102 (Kaempferol 3-O-β-D-glucopyranoside) exhibited higher average SASA values, around 9,000 Å², with noticeable fluctuations throughout the simulation. CID: 5280805 (Rutin), exhibited moderate SASA values and maintained a relatively stable profile. The lower solvent exposure of CID: 5282160 (Quercimeritrin) indicates that this complex might have a more hidden shape, which could make it more stable when binding. In conclusion, the stability metrics provided by Rg and SASA highlight CID: 5280805 (Rutin), and CID: 5282160 (Quercimeritrin) as the most promising candidates for further evaluation.

#### Molecular Surface Area (MSA) and Polar Surface Area (PSA) Analysis

The MSA of all ligand-protein complexes was analysed over a 200 ns molecular dynamics simulation (Fig. 6A). CID: 5282160 (Quercimeritrin) showed the most stable MSA profile with an average of about 7,000 Å², which means it had a tighter surface shape. The control (vincristine) showed higher MSA fluctuations, with values ranging from 7,200 Å² to 7,800 Å², suggesting greater surface flexibility and solvent exposure. CID: 5280805 (Rutin), and CID: 5282102 (Kaempferol 3-O-β-D-glucopyranoside) exhibited moderate MSA profiles, with occasional fluctuations but generally stable behaviour. These results indicate that CID: 5282160 (Quercimeritrin) forms a tighter complex than the control, which may contribute to its binding stability.

**Fig. 6:**
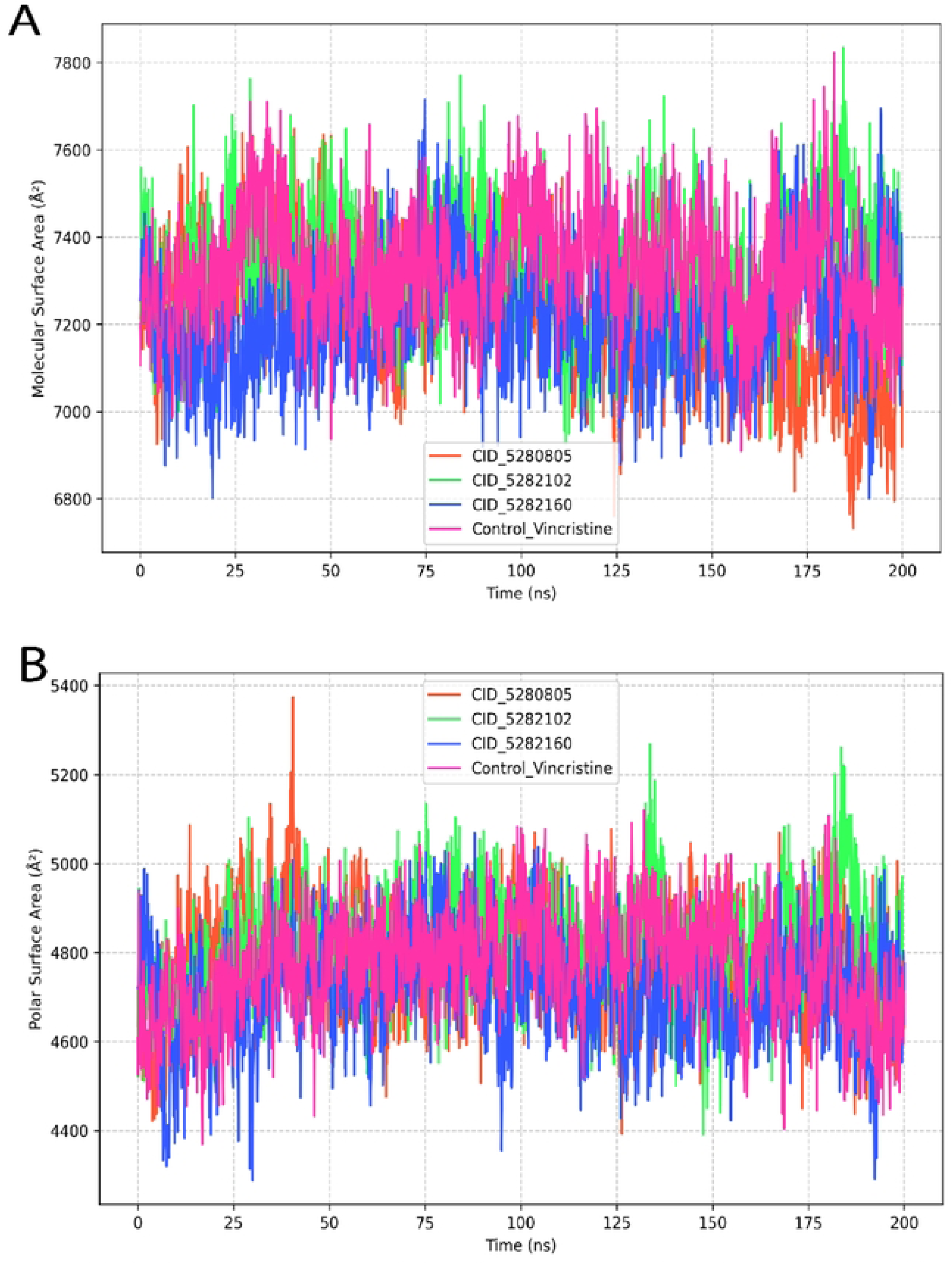
(A) Molecular surface area (MolSA) plots of protein-ligand complexes over 200 ns, indicating surface compactness (B) Polar Surface Area (PSA) plots show the solvent-exposed polar regions of the complexes.

The PSA was also evaluated to assess hydrophilic surface exposure (Fig. 6B). CID: 5282160 (Quercimeritrin) had the lowest PSA values, about 4,500 Å², which means it has very few polar groups that are exposed to the solvent. The control displayed the highest PSA values, averaging around 5,000 Å² with frequent fluctuations. CID: 5280805 (Rutin) and CID: 5282102 (Kaempferol 3-O-β-D-glucopyranoside) displayed intermediate PSA profiles, suggesting partial exposure to polar residues. The lower exposure of polar surfaces in CID: 5282160 (Quercimeritrin) might improve its ability to interact with water-repelling substances and make its binding more stable.

#### Intra-molecular Binding Pattern Analysis

The interaction fractions for each complex (Fig. 7A-7D) revealed key binding behaviors. CID: 5280805 (Rutin), mainly connected with Leu15, Glu30, and Glu128, forming strong hydrogen bonds and hydrophobic interactions. CID: 5282102 (Kaempferol 3-O-β-D-glucopyranoside) showed strong binding with Arg146 and Asn16, backed by hydrophobic and hydrogen-bond interactions. CID: 5282160 (Quercimeritrin) showed the strongest connection, with strong interactions with Ile135 and Glu128, meaning it had stable binding through both hydrophobic and polar interactions. In comparison, the control (vincristine) showed weaker and more spread-out interactions with more water involvement, indicating it is less stable.

**Fig. 7:**
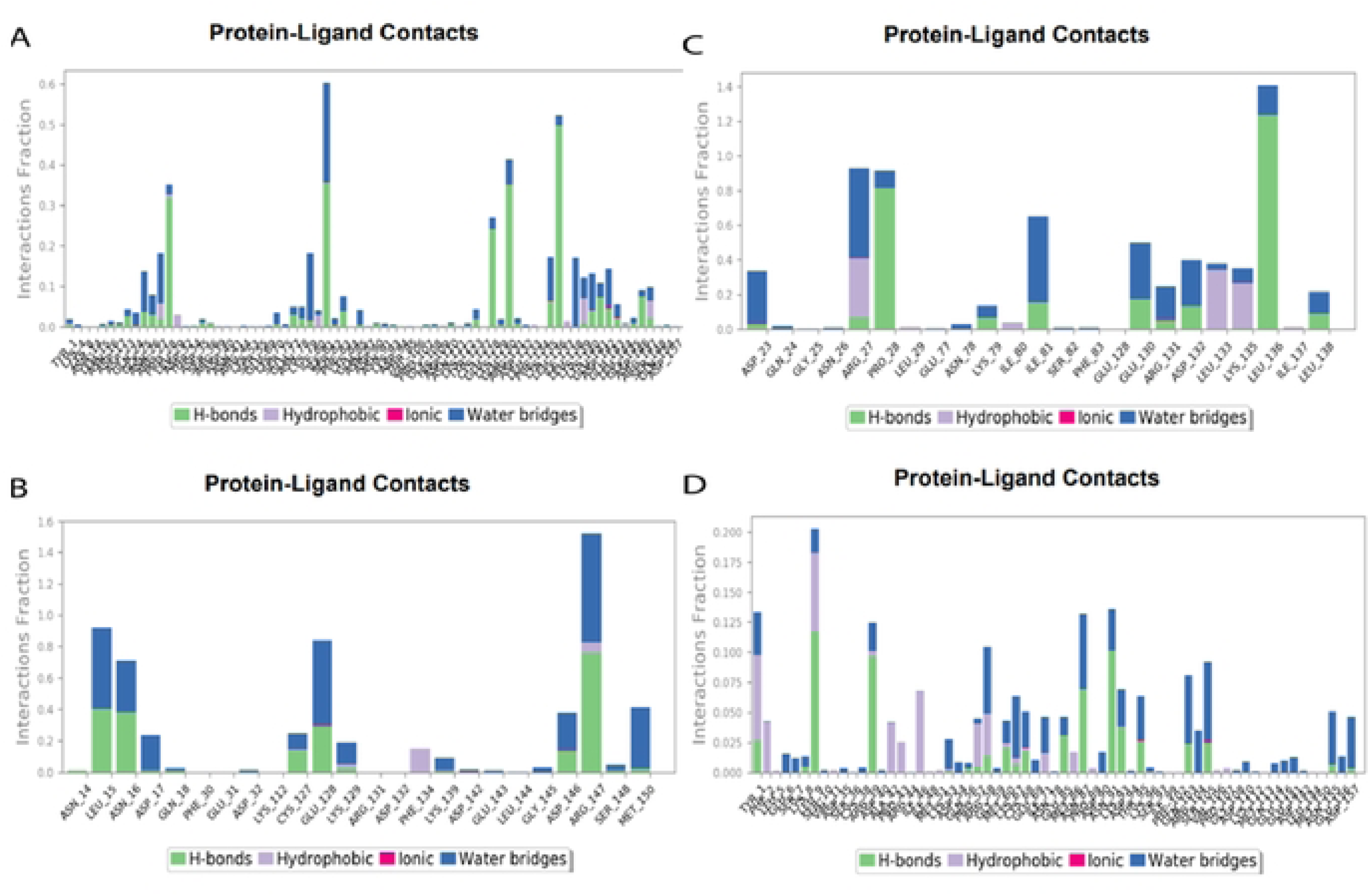
Stacked bar plots showing residue-wise interaction fractions between the ligand and protein complexes including (A) CID: 5280805, (B) CID: 5282102, (C) CID: 5282160, and (D) control ligand Vincristine. Several colours represent different interaction types, with the x-axis indicating residue positions and the y-axis showing the fraction of simulation frames with interactions.

#### MM-GBSA binding free energy analysis reveals strong ligand-receptor interactions

The MM-GBSA analysis gave us information about the binding affinities of the tested ligand-receptor complexes. CID: 5280805 (Rutin), CID: 5282102 (Kaempferol 3-O-β-D-glucopyranoside), and CID: 5282160 (Quercimeritrin) exhibited ΔG binding energies of −8.1, −7.9, and −8.3 kcal/mol, respectively, indicating favorable and stable interactions with the target receptor (Fig. 8). The control ligand (vincristine) showed a comparatively weaker binding energy of −7.5 kcal/mol. These results imply that the tested compounds, particularly CID: 5282160 (Quercimeritrin) and CID: 5280805 (Rutin), may have a higher binding potential compared to vincristine. The negative ΔG values seen in all ligands indicate that their binding is energetically favorable, suggesting these compounds could be good options for further testing in drug discovery.

**Fig. 8:**
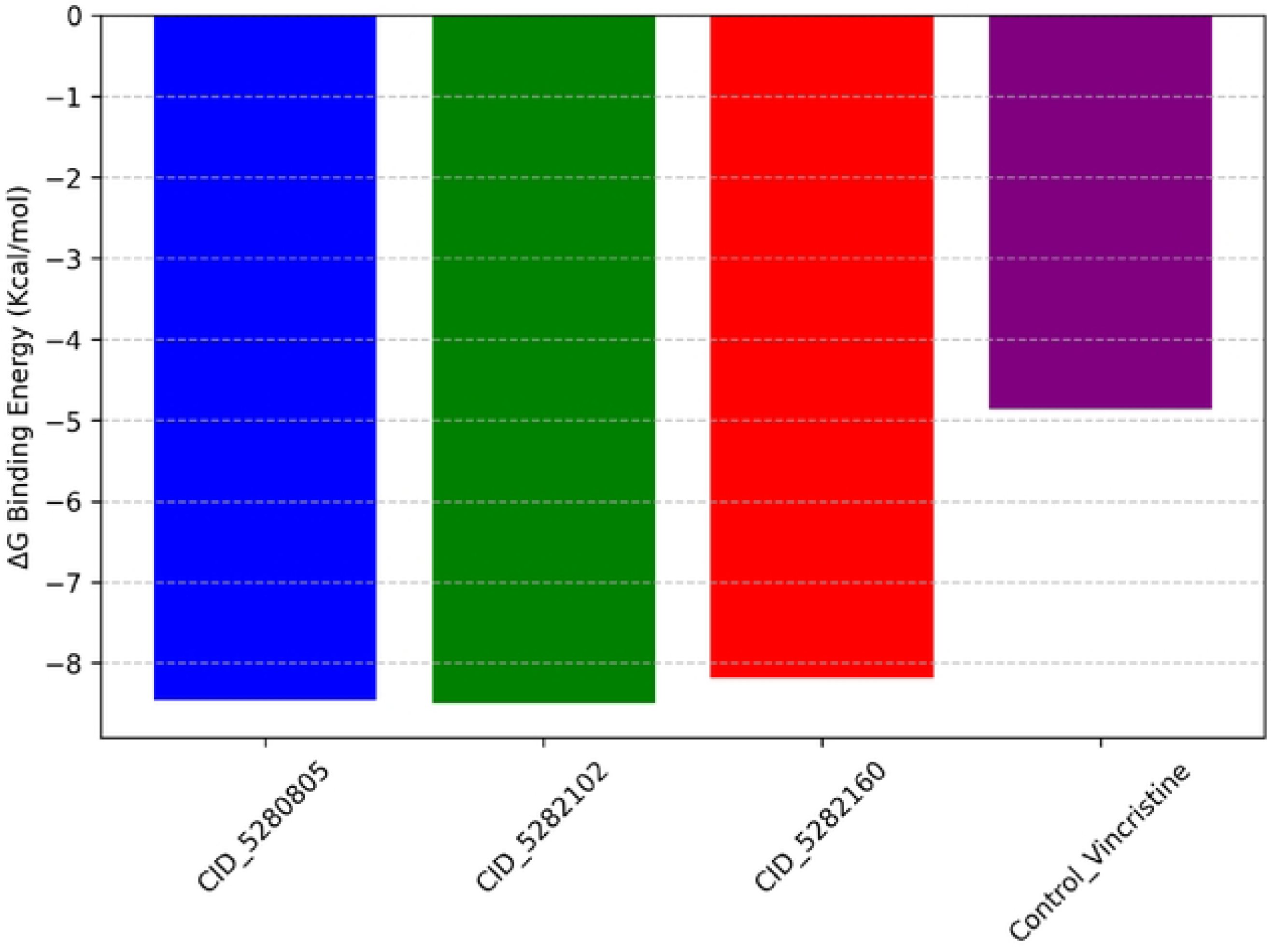
MM/GBSA-calculated binding free energies (AG, kcal/mot) for three candidate ligands (CID_528005, CID_5282102, and CID_5282160) compared to the control compound vincristine. All candidates showed more favorable binding energies than the control, indicating stronger predicted binding affinities.

## Discussion

Flora and natural substances are pivotal in medicine and serve as crucial prototypes for the innovation of new pharmaceuticals [15]. They provide a significant source of molecules exhibiting diverse biological activities and chemical structures. Certain natural compounds can specifically eradicate cancer cells while preserving healthy cells. They may achieve this by disrupting cellular proliferation, inducing apoptosis, or inhibiting angiogenesis [51]. About 60% of today’s cancer drugs came from or are inspired by natural products, showing they are effective in treating cancer. Examples include paclitaxel (Taxol) from the Pacific yew tree, vincristine from the Madagascar periwinkle, curcumin from turmeric, genistein from soybeans, and resveratrol from grapes and red wine. The present study was designed to assess the therapeutic potential of *Hylocereus polyrhizus* peel extract in the DMBA-croton oi-induced skin carcinogenic mice.

DMBA is known as indirect-acting, also organ-specific carcinogen that is metabolized by CYP1A1 and CYP1B1 enzymes of the CYP450 family into the final carcinogen 1,2-epoxide-3,4-diol DMBA [52]. DMBA intermediates induce genomic alterations by forming adducts with DNA and serve as initiators in chemical carcinogenesis [60]. 12-O-tetradecanoyl phorbol-13-acetate, the active component of croton oil, interacts with protein kinase C, inducing the transcription of early genes c-fos and c-jun, and activates the transcription factors activating protein-1 and NF-κB, culminating in cellular proliferation and death. Earlier observations indicate that administering different plant extracts during chemically induced carcinogenesis reduces the cumulative number of tumors, tumor yield, tumor burden, and tumor incidence [53, 54]. Similarly, our *in vivo* study demonstrate that the total number of tumors, tumor size, and tumor weight were decreased by the administration of HPPE compared to the carcinogenic control group. The HPPE at 500 mg/kg reduced the number of tumor formations more than 1.5 times, tumor burden more than 2 times, and tumor yield more than 2.5 times compared to the carcinogenic group, demonstrating its efficacy in skin cancer prevention (Table 2). This result may be due to the extract containing antioxidants such as phenolics and flavonoids, as early reports indicated that these compounds help prevent oxidative stress, which is associated with the onset of skin cancer [55].

Mice treated with DMBA/croton oil exhibited the reduction in the levels of total protein and endogenous antioxidant molecules in comparison to normal mice (Table 3). In comparison to the carcinogenic group, HPPE at 500 mg/kg markedly elevated the concentrations of total protein, GSH, SOD, and CAT in the tumor tissue and liver of mice, demonstrating the extract’s efficacy as an antioxidant in cancer treatment as evidenced in early studies [22, 56].

Serum biochemical parameters may provide important information about the health status of skin cancer patients and the efficacy of treatment. A variety of serum biochemical parameters (SGPT, SGOT, urea, creatinine, bilirubin, lipid profiles, etc.) can be used to evaluate the liver, kidney and heart functions which can be influenced by cancer or as a side effect of treatments such as chemotherapy [57]. In cancer patients, these biomarkers are observed to fluctuate beyond the standard range [58]. Phytocompounds have shown potential in modulating serum biochemical parameters. In comparison to the control group, the carcinogenic control group exhibited elevated serum biomarker levels (Table 4). Standard drug vincristine and also HPPE at 500 mg/kg significantly lowered these serum biomarker levels and maintained the value within the safety range. Phytocompounds such as silymarin, curcumin, quercetin etc. lowered levels of SGPT, SGOT, urea, bilirubin, creatinine, and LDL by reducing oxidative stress, inflammation, and cholesterol production [59-61]. The phenolic compounds and flavonoids present in HPPE may be responsible for this protective action for liver and kidney.

An inflammatory reaction plays an important role for expansion and promotion of cancer. Previous study indicates that boosted level of proinflammatory cytokines and inflammatory mediators play vital role in skin cancer promotion [62]. In the tumor microenvironment, these cytokines promote cancer growth and aggressiveness. Cancer can be caused by a malfunction of these molecules that control the immune system, cell growth, and inflammation. By promoting angiogenesis, inflammation, and immune suppression, TGF-β1 in skin cancer can lead to the formation of tumors [63]. It helps produce extracellular matrix proteins during fibrosis [64]. The creation of prostaglandins, pro-inflammatory mediators also promote tumor growth, angiogenesis, and metastasis [65]. COX-2 inhibition reduces tumor development and inflammation [66]. According to Park and Hong, NF-κB is a transcription factor that controls gene expression related to inflammation, cell proliferation, and survival [67]. NFκB activation in skin cancer promotes pro-inflammatory cytokines and survival genes, promoting tumor development and apoptosis resistance [64]. Constitutive NF-κB activation can cause persistent inflammation and tumor growth [68]. Inflammatory mediators often collaborate to cause skin cancer. For example, TGF-*β*1 activates NF-κB, resulting in enhanced pro-inflammatory gene expression. COX-2 can also activate NF-κB, enhancing inflammatory signaling [66]. As our in vivo result suggested that the HPPE has potential activity against skin cancer, we examined the mRNA expression levels of pro-inflammatory cytokines (TNF-*α*, IL-1β, IL-6, IL-18) and inflammatory mediators (TGF-*β*1, COX-2, NFκB) in chemically generated skin carcinogenic mice. Figure 1 shows that the mice with skin cancer had higher levels of above-mentioned genes compared to the normal mice, especially TGF-*β*1, which was more than 1.5 times higher. However, HPPE (250 mg/kg) and HPPE (500 mg/kg) significantly reduced these genes expression compared to the carcinogenic control group (fig. 1).

Flavonoids have been shown to inhibit TGF-*β* (Transforming Growth Factor Beta), a cytokine involved in various cellular processes, including cell growth, differentiation, and fibrosis [69]. By modulating TGF-*β* signaling, flavonoids can potentially reduce inflammation and fibrosis, as well as suppress tumor progression [70]. So, we attempted the molecular docking to further evaluate the interactions of reported compounds (identified flavonoids) with selected proteins. This method allows for quick testing of how proteins and ligands interact, helping to find strong binders that could lead to new treatments.

The molecular docking and binding affinity analysis found that Quercimeritrin, Rutin, and Kaempferol 3-O-β-D-glucopyranoside had binding scores of −7.4, −7.1, and −7.0 kcal/mol with the TGF-*β*1 (PDB ID: 6B8Y) protein, respectively. These values show that the binding interactions are stronger than the standard drug, vincristine (−5.3 kcal/mol), which supports the idea that flavonoid compounds usually have better binding because their structure allows for more hydrogen bonding and π-stacking interactions. Quercimeritrin demonstrated hydrogen bonding with residues Ser-148, Asn-116, and Asp-146 while forming π-stacking interactions with Phe-127, contributing to its enhanced binding stability. Similar interactions were observed for Rutin and Kaempferol 3-O-β-D-glucopyranoside, which displayed additional hydrophobic interactions. In contrast, vincristine exhibited fewer hydrogen bonds and more water-mediated contacts, indicative of weaker binding [71]. Hydrogen bonding and hydrophobic interactions are crucial in drug design, influencing binding affinity, specificity, and bioavailability. Hydrogen bonds enhance drug-target recognition and stability but may hinder membrane permeability if excessive [72]. Hydrophobic interactions improve receptor binding and membrane permeability but can reduce solubility if too strong. Achieving an optimal balance between these forces ensures effective drug absorption, distribution, and metabolism, leading to potent and selective therapeutics [73].

Drug-likeness assessment is a crucial step in identifying promising therapeutic candidates, as only a limited number of chemical compounds possess medicinal potential. Drug-likeness evaluation effectively distinguishes compounds with undesirable properties, particularly those with poor ADMET profiles, in drug discovery [74]. In our study, the ADMET analysis showed that Kaempferol 3-O-β-D-glucopyranoside had the best pharmacokinetic profile with the highest intestinal absorption rate of 54.95%. Research indicates that a compound is considered poorly absorbed if its absorption rate is below 30%, indicating limited bioavailability and potential inefficacy as a drug candidate [75]. Moderate tissue distribution and low central nervous system (CNS) permeability were observed for all compounds, aligning with previous studies indicating that glycosylated flavonoids typically have limited CNS penetration due to their polar nature. Adding sugar groups makes these compounds more water-loving, which makes it harder for them to pass through the fatty barrier of the blood-brain barrier (BBB) [76]. Lipinski’s Rule of Five serves as a key guideline in drug discovery for assessing oral bioavailability. Violating more than one of these rules suggests poor ADME properties, making the compound less suitable for oral administration [77]. In our study, even though some compounds broke Lipinski’s rules regarding molecular weight and the number of hydrogen bonds, they all had a bioavailability score of 0.17, which means they can be partially absorbed when taken by mouth. In silico toxicity testing utilizes computational models to predict the potential toxicological effects of compounds based on their molecular structure. This approach enables early identification of toxic risks, aiding in the selection of safer candidates and reducing the need for animal testing in the drug development process [78]. In our current study, we have found no AMES toxicity, hepatotoxicity, or skin sensitization risks of all reported compounds.

Molecular dynamics simulation is used to assess the stability of protein-ligand complexes, evaluating protein flexibility and stiffness under physiological conditions. RMSF values reflect the complex’s fluctuations, while RMSD analysis confirms minimal protein-ligand fluctuations, indicating complex stability [79]. In our study, MDS studies indicated greater dynamic stability for Rutin and Kaempferol 3-O-β-D-glucopyranoside against TGF-*β*1 protein, as evidenced by RMSD values stabilizing around 2.5 Å after 50 ns. A low RMSD indicates that the structure remains stable and closely aligned with the reference, while a high RMSD suggests significant conformational changes or instability in the complex [80]. Lower RMSF values for Rutin and Kaempferol 3-O-β-D-glucopyranoside reflected tighter binding and reduced conformational freedom across most residues, particularly in critical binding regions, aligning with literature suggesting that reduced flexibility is often associated with higher binding stability [81]. Additionally, Quercimeritrin showed less exposure to solvent and a more compact shape, as shown by its lower SASA and MSA values, which means it has more buried and stable binding states. Research indicates that higher SASA and MSA values indicate greater exposure and less compactness, suggesting less stability. Lower values reflect more compact structures, with tighter packing of amino acids and water molecules, indicating greater stability [82].

MM-GBSA (Molecular Mechanics Generalized Born Surface Area) analysis estimates the binding free energy of protein-ligand complexes by combining molecular mechanics, solvation models, and surface area calculations. It helps evaluate binding strength and stability, aiding in drug design and lead optimization [83]. MM-GBSA analysis further confirmed the superior binding potential of Quercimeritrin, which exhibited the most favorable ΔG binding energy of −8.3 kcal/mol, followed by Rutin (−8.1 kcal/mol) and Kaempferol 3-O-β-D-glucopyranoside (−7.9 kcal/mol). Similar studies have shown that flavonoids can exhibit strong binding free energies due to their ability to form multiple hydrogen bonds and π-stacking interactions [84]. In conclusion, Quercimeritrin, Rutin, and Kaempferol 3-O-β-D-glucopyranoside emerged as the most promising candidates for further validation.

## Conclusion

The findings of the present study suggests that *Hylocereus polyrhizus* peel extract has chemopreventive potential through upregulating endogenous antioxidants as well as suppressing different proinflammatory and inflammatory cytokines. Quercimeritrin, Rutin, and Kaempferol 3-O-β-D-glucopyranoside might be the probable leads responsible for this activity.

## Supporting information

**Supplementary Table S1:** Binding score of all reported ten compounds against corresponding ten proteins.

**Supplementary Table S2:** Binding interaction analysis of selected three ligands against TGF-*β*1 (PDB ID: 6B8Y) protein.

